# Identification of Disease Modules Using Higher-Order Network Structure

**DOI:** 10.1101/2022.12.24.521876

**Authors:** Pramesh Singh, Hannah Kuder, Anna Ritz

## Abstract

**Motivation:** Higher-order interaction patterns among proteins have the potential to reveal mechanisms behind molecular processes and diseases. While clustering methods are used to identify functional groups within molecular interaction networks, these methods largely focus on edge density and do not explicitly take into consideration higher-order interactions. Disease genes in these networks have been shown to exhibit rich higher-order structure in their vicinity, and considering these higher-order interaction patterns in network clustering have the potential to reveal new disease-associated modules.

**Results:** We propose a higher-order community detection method which identifies community structure in networks with respect to specific higher-order connectivity patterns beyond edges. Higher-order community detection on four different protein-protein interaction networks identifies biologically significant modules and disease modules that conventional edge-based clustering methods fail to discover. Higher-order clusters also identify disease modules from GWAS data, including new modules that were not discovered by top-performing approaches in a Disease Module DREAM Challenge. Our approach provides a more comprehensive view of community structure that enables us to predict new disease-gene associations.

**Availability:** https://github.com/Reed-CompBio/graphlet-clustering

## 1 Introduction

Understanding how genes and proteins interact with each other is a fundamental problem in molecular biology. Recent advancements in high-throughput experiments and computational techniques have enabled accurate inference of the underlying molecular interaction networks. Many complex diseases are caused by a number of genes or proteins interacting with one another (Oti et al., 2006; Ghiassian et al., 2015), yet identifying such a group (also called a disease module) in a molecular interaction network such as a protein-protein interaction (PPI) network (an interactome) is computationally challenging. A common way to find these groups is to use community detection (or clustering) methods that aim to find densely connected subsets of nodes in a given network. A number of different community detection algorithms have been developed and used extensively over the years for this task (Fortunato, 2010; Choobdar et al., 2019). Recently, the DREAM Disease Module Identification Challenge systematically assessed 75 community detection algorithms to detect modules across six different PPI networks that are enriched in genome-wide association study (GWAS) data from 180 diseases (Choobdar et al., 2019). While these methods have been useful in detecting disease groups in biological networks, they differ significantly from each other and show varying performance (for example, number of significant modules and their sizes), suggesting that optimal detection of these disease modules remains a challenging task.

Despite their differences, nearly all community detection algorithms focus on identifying groups of nodes that are densely connected by edges. Thus, these methods rely on pairwise relationships between nodes while neglecting higher-order interaction patterns among more than two nodes. The importance of higher-order structure within biological networks has been emphasized by many recent studies (Agrawal et al., 2018; Rubel et al., 2021; Benson et al., 2016). Therefore, it is worthwhile to extend the idea of communities to higher-order communities that are groups of nodes connected by pattern involving more than two nodes. However, besides a few methods (Arenas et al., 2008; Benson et al., 2016) which define communities of motifs (Milo *et al*., 2002), not much attention has been paid to investigating higher-order communities within networks.

Graphlets (Fig. 1) are small induced subgraphs and have been used to characterize the higher-order topology of biological networks (Pržulj *et al*., 2004; Hočevar and Demšar, 2017; Agrawal *et al*., 2018; Rubel *et al*., 2021). A recent paper defines graphlet-induced networks (Windels *et al*., 2019), in which, all nodes that comprise a graphlet in an PPI network are connected into a clique, resulting in a network with dense subregions. However, the density of the resulting network can negatively affect the quality of network-based clustering (or *partition*).

**Figure 1:**
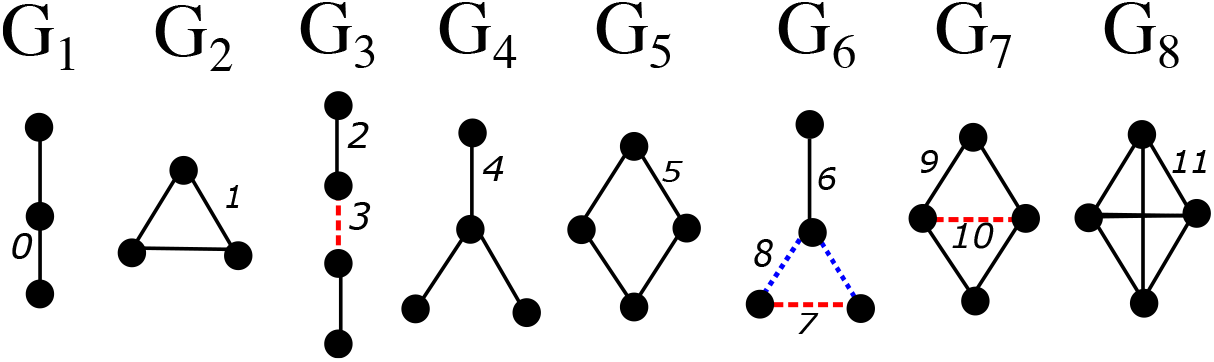
All graphlets of size 3 and 4 (*G*_1_ – *G*_8_). The distinct edge positions (0 – 11) (edge orbits) are shown with a different line style and color.

In this paper, we develop a new graphlet-based community detection method that generalizes the conventional edge-based communities and identifies groups of nodes that are connected through specific graphlets. Our approach involves transforming a given network to retain edges that participate in a particular graphlet and subsequently applying a random-walk based clustering algorithm to this transformed network. This way, by restricting the random-walk to edges of interest we can distinguish the parts of the network with a high concentration of a particular graphlet. We show that different graphlets admit quantifiably different clusterings, that graphlet enrichment depends on the density and structure of the interactome. Comparing these clusters from four different interactomes with expert curated pathway databases, we find that higher-order graphlets detect biologically relevant functional groups that are missed by the edge-based, classic clustering algorithm. Further, using GWAS trait datasets and disease association datasets, we show that specific graphlets admit clusters that are enriched for specific trait and disease-associated genes that edge-based clustering algorithms do not capture. Thus, leveraging the higher-order connectivity of networks in community detection applications can reveal relationships among disease genes that were previously unknown.

## 2 Methods

### 2.1 Graphlets

Graphlets are defined as connected, induced, non-isomorphic subgraphs of a specific size (Pržulj et al., 2004). Graphlets describe the structure of a network without requiring the specification of a null model and thus differ from motifs (Milo et al., 2002). The edges of every graphlet are partitioned into a set of automorphism groups called orbits such that two edges belong to the same orbit if they map to each other in some isomorphic projection of the graphlet onto itself (Hočevar and Demšar, 2017) (Fig. 1). There are 30 graphlets up to five nodes that have 67 edge orbits (See Supplementary Fig. S1). Existing software such as ORCA (Hočevar and Demšar, 2017) can count, for every edge, the number of edge orbits of each type.

### 2.2 Graphlet-induced community structure

To identify communities that are enriched for specific graphlets (graphlet-induced clusters or modules), we make use of the Markov Clustering algorithm (MCL) (Van Dongen, 2000) with a modified initial transition matrix.

#### 2.2.1 Markov Clustering Algorithm (Van Dongen, 2000)

Given an adjacency matrix *A*, a random walk on a network can be defined by a transition matrix *P* where the probability of transitioning from node *i* to node *j* is *p*_*ij*_ = *a*_*ij*_/ ∑_*j*_ *a*_*ij*_. To find groups of densely connected nodes (i.e. clusters) in a network, the standard MCL simulates a random walk and successively applies *expansion* and *inflation* operators on the transition matrix *P*. The expansion operation spreads random flow while the inflation operation makes strong links stronger and weak links weaker which reduces the flow between clusters. As the algorithm progresses, the network gets divided into disconnected subnetworks. The procedure is repeated until the transition matrix converges, i.e., it does not change with further expansion or inflation operations (Van Dongen, 2000). Finally, connected subgraphs that remain after convergence are extracted as clusters. The granularity of clusters can be tuned by varying the inflation parameter. In general, a larger inflation parameter results in a more fine-grained clusters.We used an existing implementation of MCL (Van Dongen, 2008). For large networks, this algorithm does not follow the MCL procedure exactly and uses approximations for speed thus we sometimes find isolated nodes in clusters. As a final step, we identify and eliminate any such nodes and retain only the connected component within the cluster.

#### 2.2.2 Modified Transition Matrix

We modify the transition matrix *P* to feed graphlet-specific transition matrices into MCL. For a given graphlet *G*_*k*_, the probability to transition from node *i* to node *j* is non-zero only when the edge from *i* to *j* participates in graphlet *G*_*k*_. Let *O*_*k*_ be the set of edge orbits that define graphlet *G*_*k*_ (e.g., *O*_*k*_ = *{*2, 3*}* for graphlet *G*_3_ in Fig. 1). Given an undirected network *G* with adjacency matrix *A*, we first count, for every edge, the number of edge orbits of each type using ORCA (Hočevar and Demšar, 2017). We then make a modified adjacency matrix *A*^(*k*)^ where

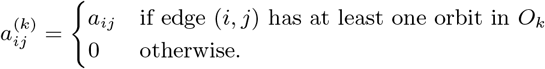

As a consequence of this edge selection, a subgraph of the original network is retained. Further, the graphlet-specific matrix for *G*_0_ is simply the original adjacency matrix *A*. The graphlet-specific transition matrix *P*^(*k*)^ for graphlet *G*^*k*^ is then

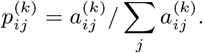

For nodes *i* that have no incident edges we set 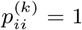, implying that a walker starting at node *i* stays at node *i*. The idea is illustrated in Fig. 2 for graphlet *G*_2_ (triangle).

**Figure 2:**
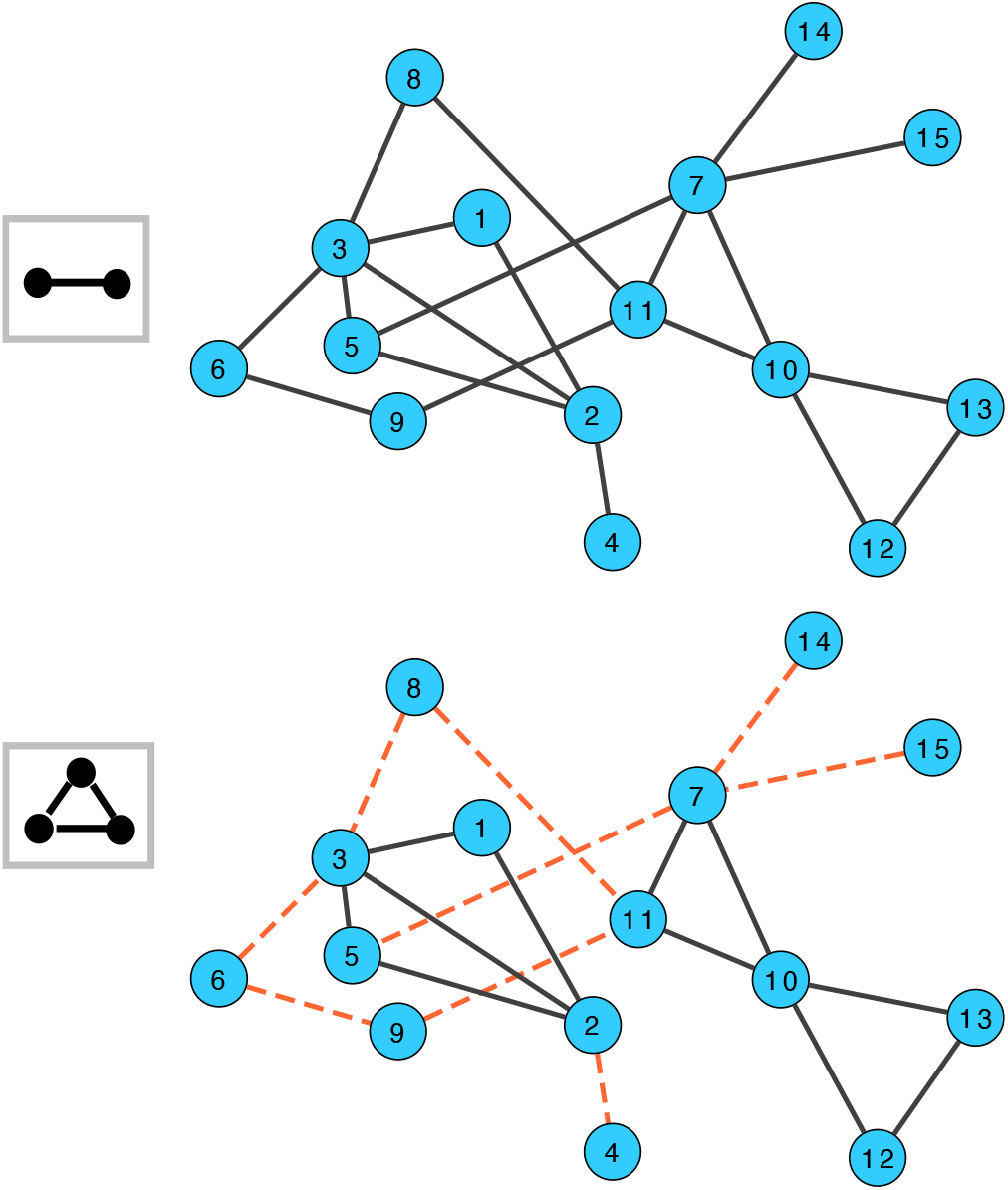
An illustration of graphlet-induced network for *G*_2_. Under the standard MCL, transition all edges are allowed and are shown by black lines (top). For graphlet *G*_2_ (triangle), red dashed edges represent transitions that are no longer allowed for *G*_2_ (bottom).

MCL is then used to find communities using the graphlet-specific transition matrices, which finds the communities that are connected through *G*_*k*_. We only keep clusters of size 3 or larger for further analysis. For the remainder of this paper, we will refer to the transformed network obtained by a modified transition network as a *graphlet-induced network*.

### 2.3 Retaining Non-Redundant Graphlet-Induced Networks

Our approach returns 30 different clusterings of the same PPI network, one for each of the graphlets up to five nodes (*G*_0_ *−G*_29_). However, some of graphlet-specific transition matrices *P* ^(*k*)^ are in fact not very different from the original transition matrix *P*. As a result, some of the MCL clusterings are similar not because they clustered different networks in the same way, but because they are essentially clustering the same network. We identify and ignore these redundant graphlets that do not alter the network. For each network, we retain graphlets *G*_*k*_ where the graphlet-induced adjacency matrix is less than 95% similar, indicating that more than 5% of the edges are dropped because they do not participate in the same graphlet (Supplementary Fig. S3). The numbers of retained graphlets for each network are shown in Table 1.

**Table 1:** PPI networks used in this study. Retained graphlets are those whose graphlet-induced transition matrices are sufficiently different from the original (*G*_0_) network. For each network, the selected inflation parameter, the size of the largest cluster, and the total number of clusters that contain at least 3 nodes is shown.

| Interactome | Network Statistics |  |  |  | Retained Graphlets | Graphlet-MCL Clustering |  |  |
| --- | --- | --- | --- | --- | --- | --- | --- | --- |
|  | Nodes | Edges | Avg. Deg. | Weighted |  | Inflation | Largest cluster | #Clusters |
| STRING (Szkarczyk et al., 2021)<br>(weights > 0.8) | 10375 | 213996 | 41.2 | yes | 1,3,4,5,7,9,10,11<br>12,15,16,17,19,20,21<br>22,24,25,27,28,29 | 8.0 | 798 | 917 |
| InWeb (Li et al., 2017) | 12420 | 397309 | 64.0 | yes | 5,8,20,22,24<br>26,27,28,29 | 3.0 | 99 | 1354 |
| SNAP (Agrawal et al., 2018) | 21557 | 338636 | 31.4 | no | 2,7,8,18,22<br>24,26,27,28,29 | 3.0 | 572 | 1735 |
| HuRI (Luck et al., 2020) | 9094 | 63242 | 14.0 | no | 2,5,7,8,18,20<br>21,22,23,24,25<br>26,27,28,29 | 3.0 | 131 | 1013 |

### 2.4 Methods for Comparison

We note that there are many available clustering algorithms, and many of them can be adapted to use graphlet-induced networks as described above. Running MCL with *G*_0_ is equivalent to the original MCL algorithm, since the *G*_0_-induced subnetwork of *G* is simply *G*. Thus, we use the *G*_0_ MCL as a comparison to traditional MCL.

We also compare our results to six additional methods from two bodies of work In the first method we consider a different way of constructing the graphlet-induced network described in Windels et al. (2019), which has similar goals to our work. In contrast to our approach, the approach of Windels et al. (2019) defines two nodes to be adjacent in a graphlet-induced network if they share a graphlet regardless of whether or not they are connected in the original network. Thus, the main difference between the two approaches is that Windels et al. (2019) transforms the network by turning a graphlet into a fully-connected clique which makes the induced networks denser than the original whereas we remove edges in a targeted manner which results in a sparser induced network than the original. To evaluate the influence of the graphlet-transformed networks, we run MCL on the transformed networks for both approaches. Due to the density of the transformed network using the graphlet-induced network of Windels et al. (2019), we limited our comparison to graphlets up to four nodes.

Next, we compare our approach to the five top-performing DREAM challenge algorithms (Choobdar et al., 2019), which include a kernel clustering approach (method K1), a random-walk based method (method R1), a local agglomerative clustering (method L1), and two methods optimizing modularity (methods M1 and M2). When comparing using the DREAM challenge algorithms, we evaluate the communities based on the DREAM Challenge inputs of 180 GWAS datasets. We note that we have not optimized our graphlet-induced MCL to perform well for the DREAM Challenge inputs, but this provides a nice baseline compared to the state-of-the-art.

### 2.5 Data Sources

#### 2.5.1 Interactomes

We applied our method to four interactomes (Table 1): InWeb (Li et al., 2017), an interactome from the Stanford Network Analysis Project (SNAP) (Agrawal et al., 2018), HuRI (Luck et al., 2020), and a subset of the STRING (Szklarczyk et al., 2021) network of edges weighted 0.8 or larger on a scale from 0 to 1. These interactomes range in size from 63, 000 edges to nearly 400, 000 edges. Given such variability in network structure, a single choice of inflation parameter may not be appropriate for all the interactomes. Thus, we performed a parameter sweep for each interactome by running MCL with varying inflation parameters (between 1.0 and 8.0) and plotted both the number of clusters returned as well as the number of nodes in the largest cluster for each network and inflation parameter (Supplementary Fig. S2). The inflation parameter was chosen such that the size of the largest cluster is of the order of hundreds of nodes (Table 1).

#### 2.5.2 Biological Process and Disease Gene Sets

To assess the performance of different graphlet-induced modules, we compare them to pathways (represented as gene sets) from the Human Molecular Signatures Database (MSigDB) (Subramanian et al., 2005). Specifically, we considered a collection of 292 pathway gene sets from BioCarta (Nishimura, 2001), 186 pathway gene sets from the Kyoto Encyclopedia of Genes and Genomes (KEGG) (Kanehisa and Goto, 2000), and 196 pathway gene sets from the Pathway Interaction Database (PID) (Schaefer et al., 2009). Curated by domain experts, these gene sets are canonical representations of a biological process.

We also compared the graphlet-induced modules with disease gene sets. We used 519 annotated disease gene sets (Agrawal et al., 2018) from DisGeNET (Piñero et al., 2015) which integrates expert-curated databases that cover information on Mendelian and complex diseases and 34 gene-level cancer datasets from The Cancer Genome Atlas (TCGA) mutations, curated by OncoVar (Wang et al., 2020).

#### 2.5.3 GWAS Trait Datasets

We also evaluated disease-gene associations using GWAS data, which offer a complementary perspective to the disease gene sets. We used a collection of 180 Genome-Wide Association Studies (GWAS) Datasets of disease-related human phenotypes from the DREAM challenge (Choobdar et al., 2019), which cover a wide range of molecular processes.

### 2.6 Module Assessment

#### 2.6.1 Evaluating Clustering Similarity

We first evaluate the similarity of MCL clusterings from the same PPI network using different graphlet-induced networks. To compare clusterings from different graphlet-induced MCL runs, as well as compare graphlet-induced MCL to the other approaches, we use the Adjusted Rand index (ARI) which controls for cluster matching due to random chance. In computing the ARI, we only include the clusters that have at least three nodes.

#### 2.6.2 Hypergeometric-Based Enrichment

To evaluate the enrichment of the pathway and disease gene sets, we use measures based on the hyper-geometric *p*-value. For every module/gene set pair, we calculate the hypergeometric *p*-value adjusted by the Benjamini-Hochberg method (Benjamini and Hochberg, 1995) for multiple hypothesis testing. We then calculate the following measures using a *p*-value cutoff of *<* 0.05:

1. The *gene set coverage* is the number of gene sets for which a significant module was found. We also calculate the *gene set percentage* as the fraction of genes (out of all genes in the gene sets) in the significant gene sets.
2. The *module coverage* is the number of modules that are significantly enriched for some gene set. We also calculate the *module percentage* as the fraction of genes in the significantly enriched modules (out of genes in all modules within the specified size).

#### 2.6.3 DREAM Challenge Enrichment

To evaluate the enrichment of genome-wide association studies (GWAS) datasets, we use the DREAM Challenge’s framework of using *Pascal* (Lamparter *et al*., 2016). Pascal first obtains gene scores by aggregating single nucleotide polymorphism *p*-values from GWAS, while correcting for linkage disequilibrium structure. It then combines the scores of genes that belong to the same pathways to obtain pathway scores (*p*-values) using the chi-squared method described in Lamparter et al. (2016). These pathway level *p*-values are further adjusted for multiple testing using the Benjamini-Hochberg correction (Choobdar et al., 2019). Finally, these adjusted *p*-values are used to determine the number of significant modules for a given method. Note that the DREAM Challenge’s criteria of number of significant modules is equivalent to our measure of *module coverage* described above. In the DREAM challenge methods are ranked according to the number of significant modules that they discover. Since each significant module can associated with more than one GWAS, we also report the number of significant GWAS datasets (*trait coverage*, similar to gene set coverage in the previous section).

## 3 Results

### 3.1 Graphlets admit different clusters

We first quantify the similarity in the community structure found by clustering different non-redundant graphlet-induced networks with MCL. The variability in the ARI scores between different MCL clusterings for each of the four PPI networks shows that a number of these clusterings are substantially different from the original *G*_0_-based clustering and thus contain distinct information within them (Fig. 3 [See Supplementary Fig. S4 for all 30 graphlets]). Specifically, about 57% pairs of community structures in STRING, 44% in InWeb, 26% in SNAP, and 58% in HuRI have an ARI score of less than 0.8.

**Figure 3:**
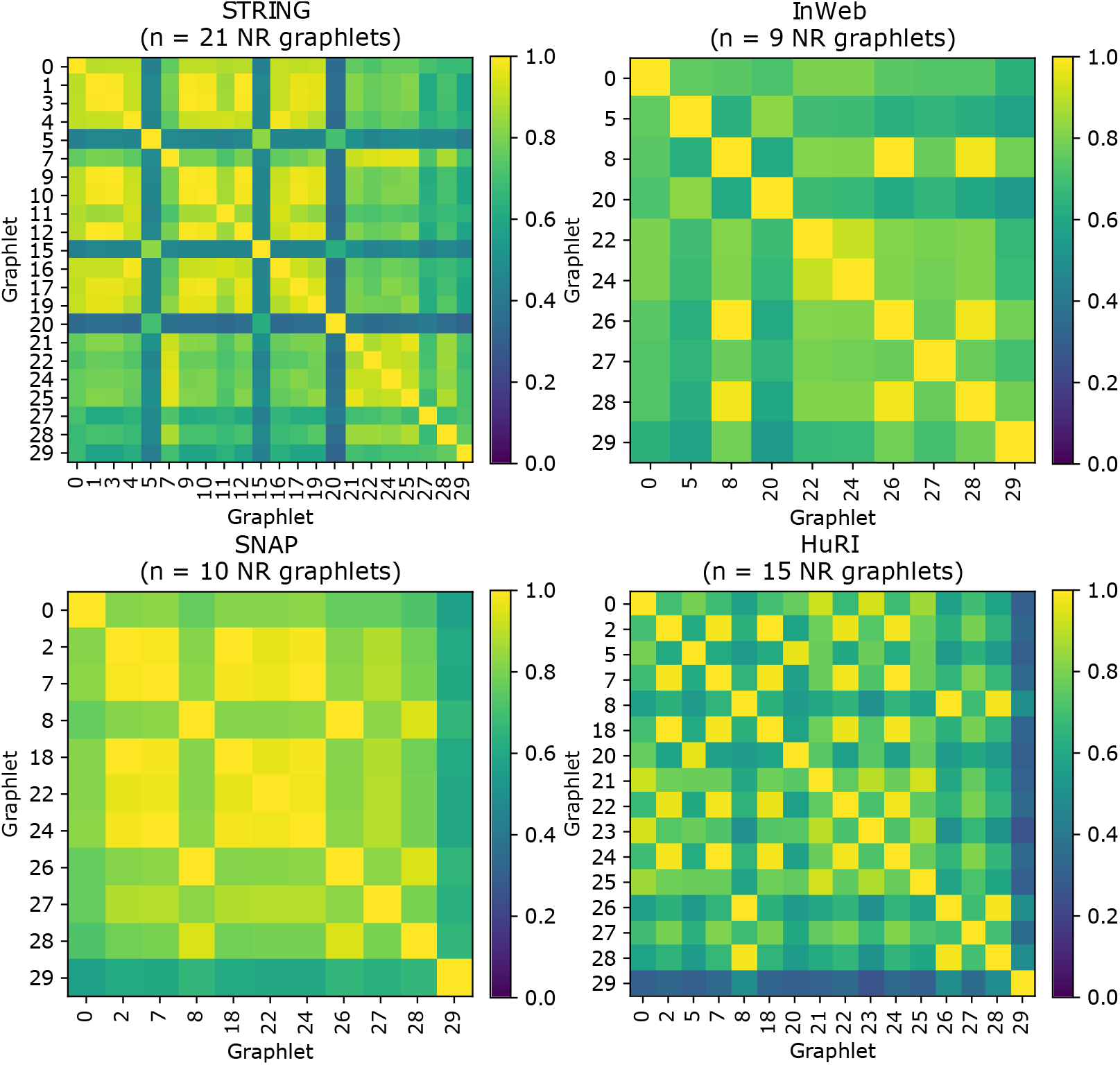
ARI scores for non-redundant (NR) graphlet-induced clustering of the interactomes. A higher ARI indicates a higher level of similarity between the two clusterings.

### 3.2 Pathway enrichment of clusters

We next assess the biological relevance of clusters obtained by higher-order graphlets. Since we expect genes within pathways to be near each other in PPI networks, we focus on the gene set percentage of the three pathway databases (BioCarta, KEGG, and PID).

In all combinations of PPI network & pathway database, the gene set percentage is larger for higher-order graphlet-induced clusterings (*G*_*k*_ = *G*_1_ *−G*_29_) than clustering based on *G*_0_ (Fig. 4). This is even true for the HuRI PPI network, which also identifies a good number of pathway-enriched modules using *G*_0_ that are not found in the higher-order clusterings (red bars). Importantly, this analysis shows that higher-order graphlets can find unique pathway associations (orange bars) that are not detected by *G*_0_.

**Figure 4:**
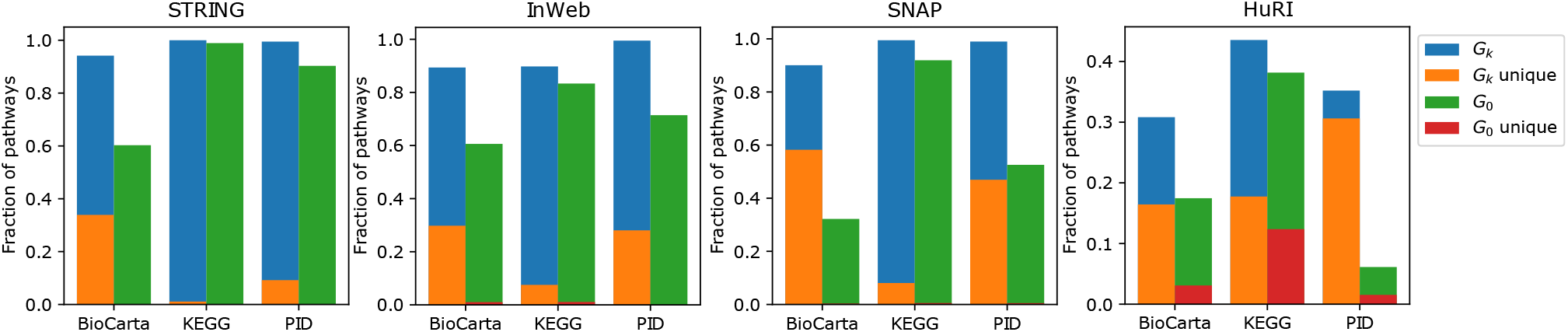
Fraction of pathways from different databases that are significantly associated with modules discovered by high-order clustering. It shows the fractional coverage obtained by *G*_0_ (green) and other graphlets *G*_*k*_ (blue) for each interactome. The height of red and orange bars shows what fraction of these pathways are unique to the two sets *G*_0_ and *G*_*k*_ respectively.

### 3.3 Disease gene enrichment

We then moved on to assessing the graphlet-enriched clusters with respect to identifying disease modules. We do not expect genes from each disease to be near each other in the PPI networks, especially for complex diseases – thus, we considered the total number of diseases that are enriched (gene set coverage) rather than the percentage of diseases found for each dataset. We find the number of significant disease modules for *G*_0_ and higher-order *G*_*k*_-based clusterings (Fig. 5). Each of these sets of clusterings finds unique disease associations and with the exception of HuRI, the number of unique associations in higher-order clusterings is consistently higher across all interactomes and disease databases. These results indicate that modules found by other graphlet-based methods can provide a large number of new disease associations that the traditional approach does not find.

**Figure 5:**
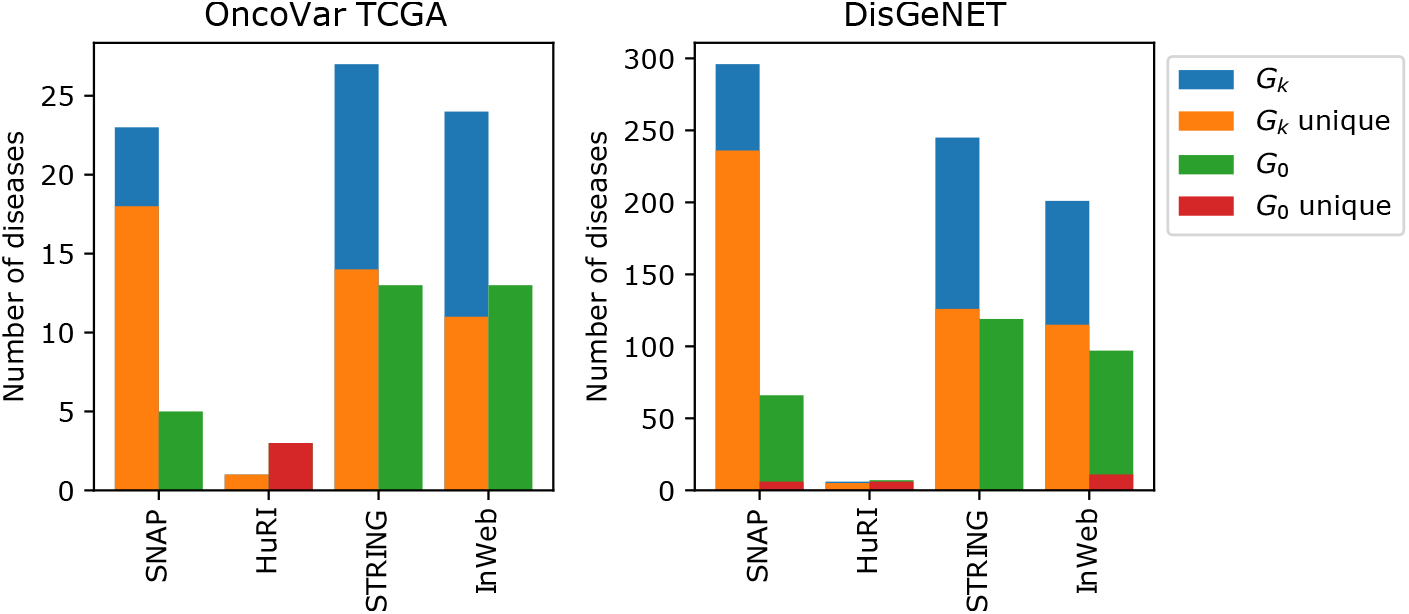
Number of significantly associated diseases from different disease association databases discovered by higher-order clustering.

To show that these modules can provide functional predictions for un-annotated genes, we examined four modules that are enriched in distinct diseases from DisGeNET in the SNAP PPI network (Fig. 6). Notably, none of these diseases are revealed in the *G*_0_-based clustering. Even the best corresponding modules in the *G*_0_-based clustering have adjusted *p*-values that do not cross the significance threshold (Supplementary Table S1). These networks show the edges that participate in the corresponding graphlet in the SNAP PPI network.

**Figure 6:**
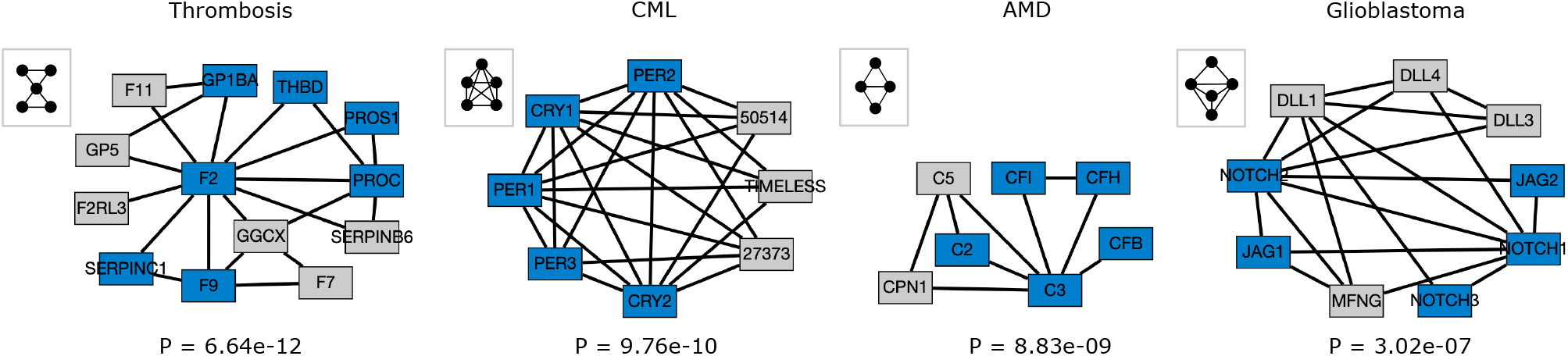
Disease modules discovered by graphlet-aware community detection using specific graphlets for Thrombosis (*G*_18_), Chronic Myeloid Leukemia (*G*_29_), Age related macular degeneration (*G*_7_), and Glioblastoma (*G*_26_). Blue nodes indicate genes present in the DisGeNet disease set and gray nodes are not annotated to the disease. The hypergeometric *p*-value is indicated below each module.

Fig. 6(a) is associated with thrombosis – a condition characterized by formation of clots inside blood vessels. The genes colored blue are already associated with thrombosis according to DisGeNet. Five of the gray genes (*GGCX, F2RL3, F11, F7*, and *GP5*) are known to play a role in blood coagulation (Megy et al., 2019). However, the gene *SERPINB6* does not have a known link to thrombosis and it may be a promising candidate for further investigation. Similarly, the association of the blue nodes with Chronic Myeloid Leukemia (Fig. 6(b)) is already known (Sheng-Fung et al., 2004; Hanoun et al., 2012), while the gray genes are potentially new. In Fig. 6(c), genes *C5* and *CPN1* are our predicted associations for age related macular degeneration besides the known involvement of the blue genes (Lu et al., 2018). Fig. 6(d) shows the disease module associated with Glioblastoma. Deregulation of NOTCH receptors and their ligands (nodes *NOTCH1, NOTCH2, NOTCH3, JAG1, JAG 2, DLL1, DLL3*, and *DLL4*) are known to play a role in Glioblastoma (Fiaschetti et al., 2014); but the role of *MFNG* is not established.

We also compared the graphlet-induced clusters from the SNAP PPI network to those using the approach of Windels et al. (2019) for graphlets up to four nodes. For the disease gene sets from DisGeNET (Piñero et al., 2015), we find that our approach results in a larger number of enriched disease modules for each graphlet *G*_1_ *− G*_7_ (Supplementary Fig. S5). Running the clustering algorithm was prohibitively slow for the other graphlet-induced networks constructed according to Windels et al. (2019), limiting our ability to compare across all four networks.

### 3.4 GWAS Enrichment

We evaluate the performance of our method to find disease/trait associated modules in the InWeb interactome using 180 GWAS datasets from the DREAM Challenge (Choobdar et al., 2019). We chose to use InWeb for this analysis since it was also used in the DREAM Challenge. InWeb is also sparse enough and can be efficiently analyzed without the need to discard the low weight edges for speed as we did in STRING (Table 1).

Our method is able to capture a number of trait-associated modules in InWeb (Fig. 7). While none of the graphlet-based methods outperform K1, M1 and M2, five of these methods are at least as good as L2 and R1 (Fig. 7 (top)). We observe a similar trend we comparing the number of GWAS associations discovered by each of these methods. DREAM challenge methods K1 and M2 still find more GWAS associations, which is not necessarily surprising since the DREAM challenge methods are customized to perform well on the challenge datasets. Nevertheless, all graphlet-based methods are at least as good as the L2 (showing lowest performance of the top 5 methods) (Fig. 7 (bottom)). Interestingly, the modules found by each higher-order graphlet reveal 17 significant associations that are not found by *G*_0_ (shown in green in Fig. 7). In addition, we also find that there are five unique associations that higher-order graphlets identify but are not found by *G*_0_ or any of the five top-performing DREAM methods (Table 2). These five datasets represent four distinct disease classes, including anthropometric, cardiovascular, glycemic, and neurodegenerative diseases (Supplementary Table S2).

**Table 2:** GWAS traits identified by different higher-order graphlet-based clusterings that are not identified either by our *G*_0_-based method or any of the top 5 DREAM challenge methods.

| GWAS Trait | Graphlet<br>MCL | Module<br>Size | Adj.<br>$p$-value |
| --- | --- | --- | --- |
| Coronary Artery Disease<br>(Nikpay et al., 2015) | 22 | 3 | 9.78e-6 |
| Body Mass Index (Horikoshi et al., 2015) | 29 | 13 | 1.01e-4 |
| Type 2 Diabetes (Morris et al., 2012) | 27 | 6 | 7.86e-5 |
| Overweight (Berndt et al., 2013) | 24 | 5 | 4.02e-5 |
| Alzheimer's Disease<br>(Lambert et al., 2013) | 5 | 3 | 9.41e-6 |

**Figure 7:**
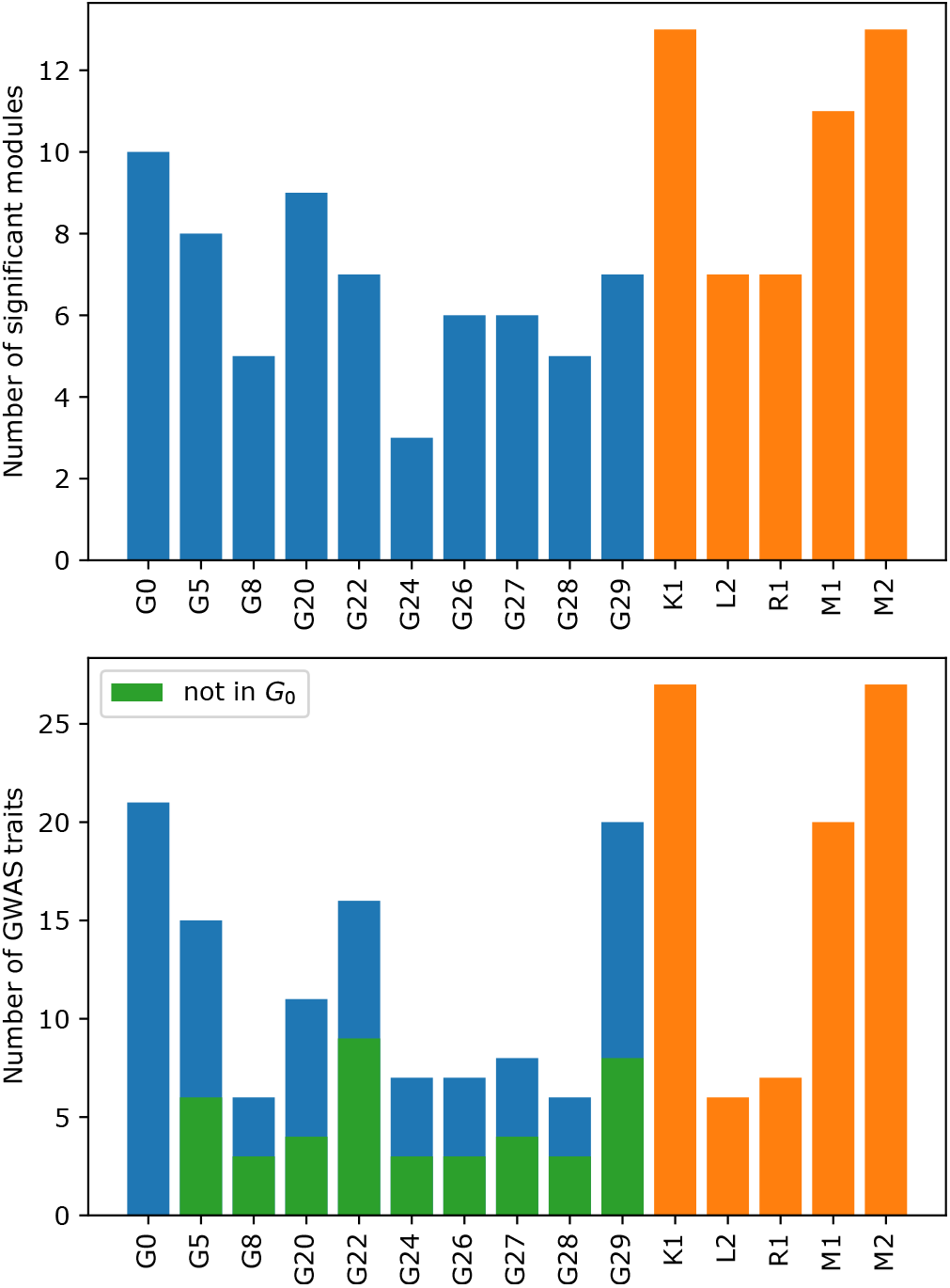
Number of modules significantly associated a GWAS trait (top) and the number of significantly associated GWAS traits found (bottom) in the InWeb interactome using different (non-redundant) graphlet-based clustering. Results obtained by different (non-redundant) graphlet-based clustering are shown in blue and top five methods from DREAM challenge submission are shown in orange whereas green indicates the set of unique associations identified by higher-order graphlets that are not found by either *G*_0_.

Two notable examples of a module uniquely identified by graphlet-based method is shown in Fig. 8 (The other three modules are visualized in Supplementary Fig. S6). First, the module of size 13 associated with body mass index (BMI) contains genes with statistically significant *Pascal* gene scores (adjusted *p* < 0.05). The involvement of genes in obesity/BMI is supported by other studies. A recent study links variants in *TAOK2* to human obseity (Agrawal et al., 2021). Genes *MAP2K3* and *MAPKAPK3* are found to play a role in BMI (Bian et al., 2013; Shao et al., 2022). Other genes in MAPK signaling that are not significant according to the gene scores (*RPS6KA4, DUSP4, MAPKAPK5*) have also been associated with obesity (Ow and Kuznetsov, 2015). *EEF2K* is also predicted to be a novel target for obesity (Joshi et al., 2021). However, this module also contains genes (e.g., *ELK1, MAPKAPK2*) with no gene scores that could potentially be associated with BMI.

**Figure 8:**
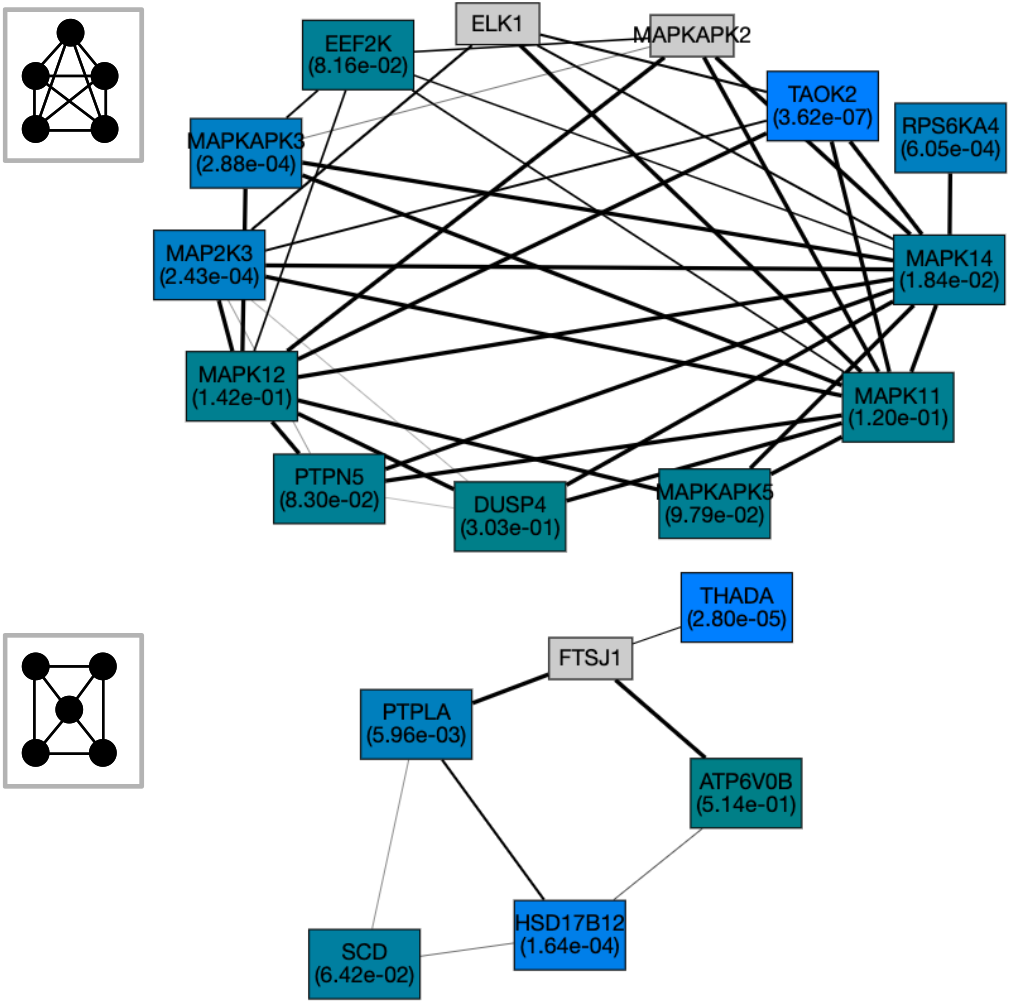
Modules detected by *G*_29_-based and *G*_27_-based community detection which are associated with the traits BMI (top) and Type 2 Diabetes (bottom) with module *p*-values 1.01e-4 and 7.86e-5 respectively computed by *Pascal* (Lamparter *et al*., 2016) (See Supplementary Table S2). The gene *p*-values in each module are indicated by different colors.

The module related to Type 2 Diabetes contains three genes with significant *p*-values (*THADA, PTPLA, HSD17B12*) and three with *p* > 0.05 (Fig. 8). The genes *THADA* and *HSD17B12* have previously been implicated in type 2 diabetes (Zeggini et al., 2008; Hachim et al., 2020). The other genes in the module are potentially new and could be good candidates for further investigations. This includes the genes *PTPLA* with a statistically significant gene score and *FTSJ* with no assigned gene score.

## 4 Discussion

Well-established community detection methods for disease module prediction often neglect the higher-order connectivity patterns among genes or proteins of interest and focus only on their pairwise relationships, even though, studies have demonstrated that higher-order structure is biologically relevant. In this paper, we have presented a generalized community detection method that incorporates higher-order structures in the form of graphlets. By providing an ensemble of partitions, each corresponding to a different graphlet (including *G*_0_), our approach provides a comprehensive view of network communities. Each of the nonredundant graphlet-based clusterings offers a unique perspective and thus, compliments the traditional (*G*_0_) clustering method. Using a collection of diverse expert-curated association datasets (pathways, diseases, and GWAS), we further demonstrated usefulness of our approach in identifying disease modules in four different interactomes. Our analysis shows strong evidence that the higher-order graphlet-based clusters can reveal unique biological associations that traditional methods can not.

A limitation of our study is that although it can find potentially novel associations using an ensemble of graphlet-based clusters, it does not determine the role or provide an interpretation of a graphlet structure in a specific biological context. While some network motifs (e.g., feed forward loop) and their functions are well-studied, a clear biological interpretation of a general graphlet structure is missing and presents a promising direction for future research.

Our graphlet-based clustering framework uses undirected graphlets and thus works with undirected networks. However, many gene/protein interactions are inherently directed. If we ignore the edge directions, we can still apply our framework to find relevant modules in these networks. However, by neglecting the directionality of edges, we may be missing important context about these interactions. An interesting future research direction will be extend this framework to directed networks by incorporating directed graphlets (Sarajli’sc et al., 2016; Trpevski et al., 2016) into our module detection approach.

Complex diseases likely have multiple factors in play, and while the same disease might appear in multiple modules, in some cases, it is possible that one graphlet is insufficient to capture the heterogeneity of disease genes. Prior work has suggested that multiple graphlets are over-represented in the same disease module or pathway (Agrawal et al., 2018; Rubel et al., 2021). Thus, extending our framework to identify modules with respect to a combination of different graphlets may reveal more accurate disease associations.

Finally, it will be interesting to investigate higher-order organization in problems beyond network clustering. For example, pathway reconstruction where the goal is find a subnetwork that connects genes of interest (Ritz et al., 2016) or the problem of detecting active modules given a set of active/seed genes (Levi et al., 2021). Even though these networks contain higher-order structure within them (Rubel et al., 2021), to our knowledge no method explicitly focuses on optimizing the higher-order structure in these networks. Thus, it may be worthwhile to develop new methods that aim to identify subnetworks that exhibit a desired graphlet profile.

## Acknowledgements

This work was supported by the National Science Foundation (NSF-DBI-1750981) to AR. We would like to thank Anthony Gitter, Lixing Yi, Max Bennett, Henry Jacques, Larry Zeng, Nina Young, and Alex Richter for fruitful discussions.

