## Supplementary Material for "Identification of Disease Modules Using Higher-Order Network Structure"


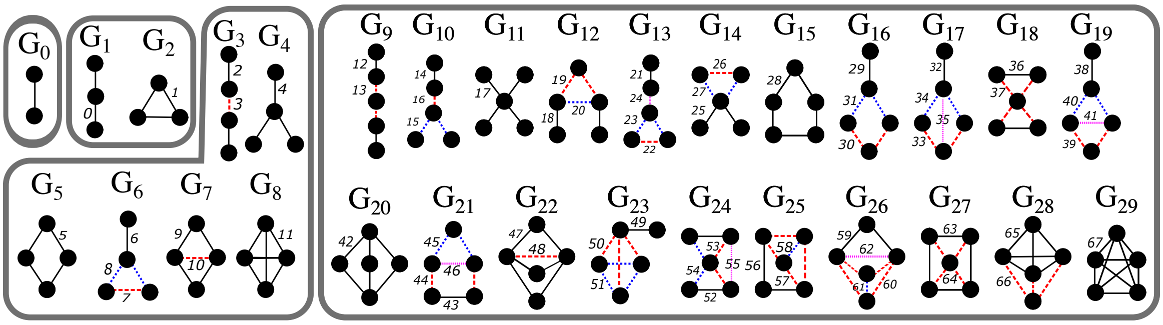

Figure S1: All 30 graphlets up to five nodes with 68 $(0-67)$ edge orbits indicated by different style and colors. Graphlet $G_0$ is a simple edge and does not have an edge-orbit label.

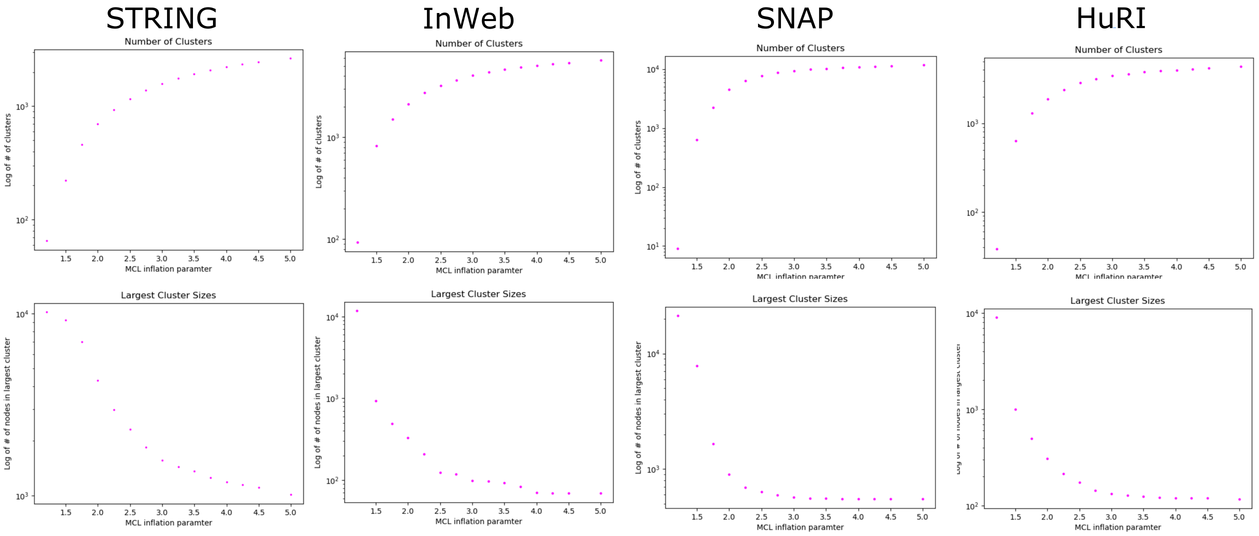


Figure S2: Parameter sweep plots for STRING, InWeb, SNAP, and HuRI. MCL was performed on each interactome's $G_0$ network for varying inflation values. The first plot for each interactome shows the log of the number of clusters plotted against the inflation parameter used. The second plot for each interactome shows the log of the number of nodes in the largest cluster plotted against the inflation parameter used.

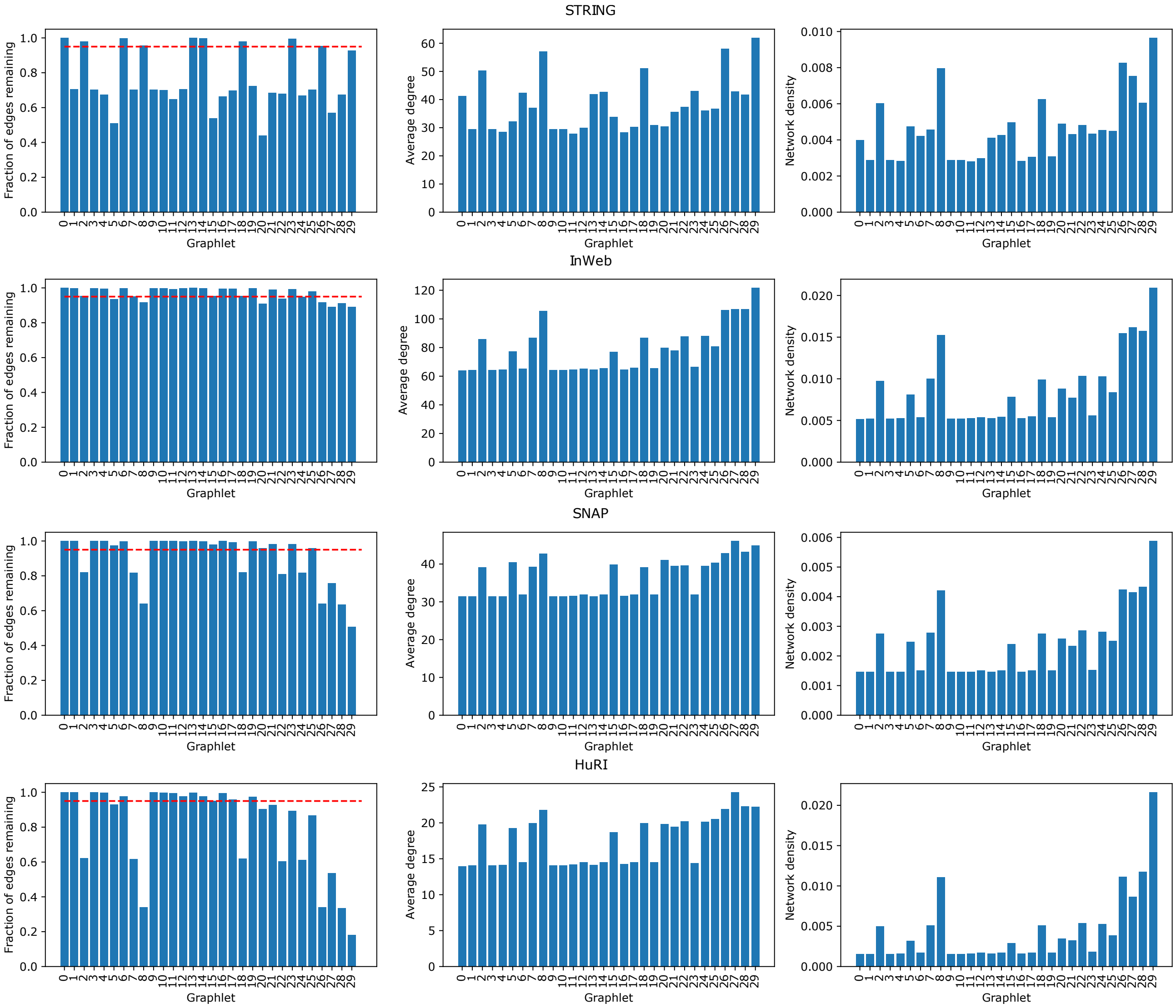


Figure S3: Global properties of modified networks corresponding to graphlets $G_0 - G_{29}$ for all four interactomes considered. The red dashed line represents fraction of edges $= 0.95$ of the original $(G_0)$ network. Networks with the remaining fraction of edges larger than this threshold are considered redundant.

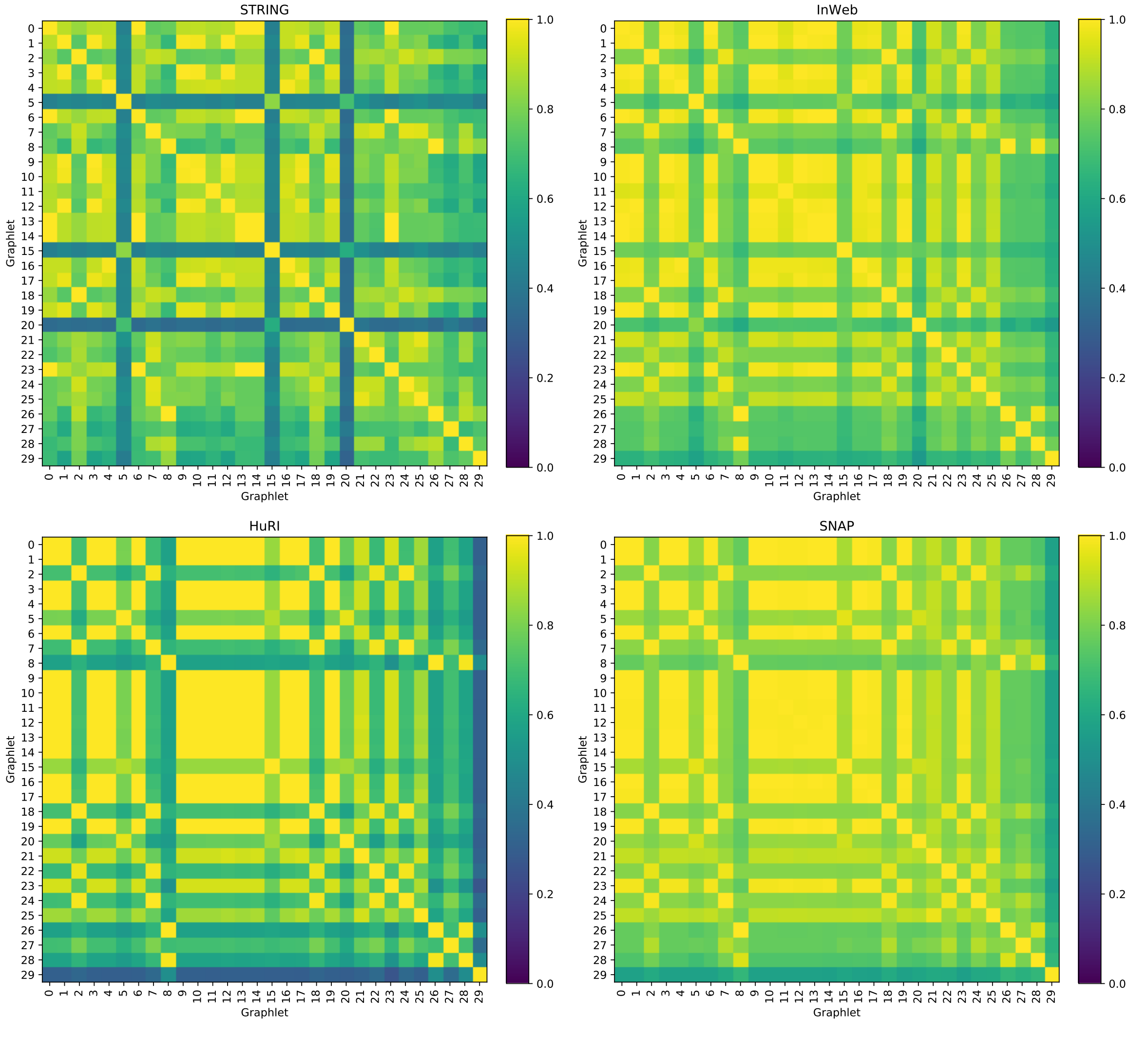


Figure S4: ARI between different graphlets-based clusterings (with respect to all graphlets $G_0 - G_{29}$).

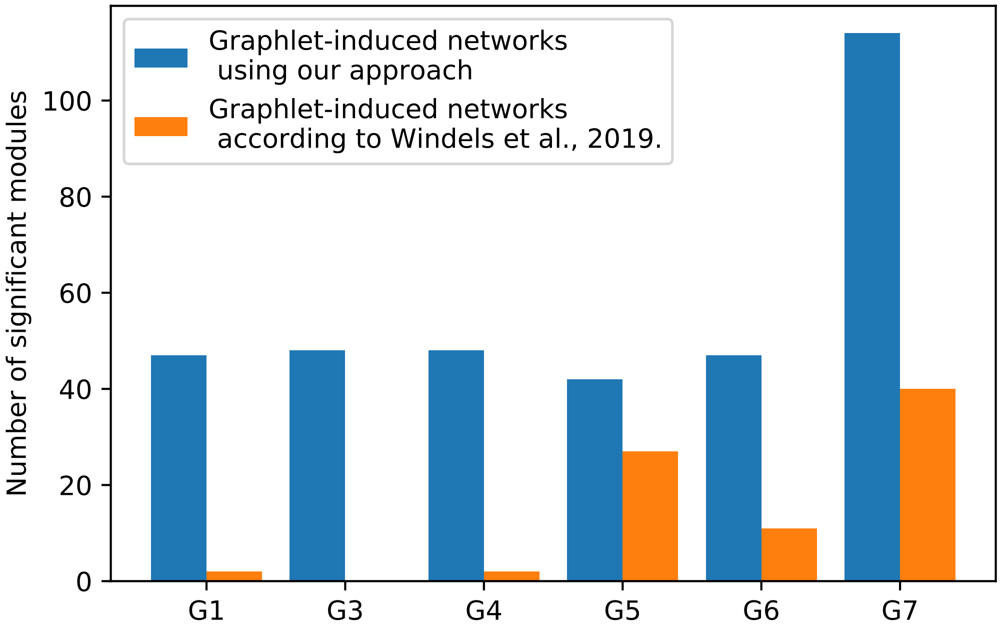


Figure S5: Number of significant modules in two different graphlet-based module detection methods in the SNAP interactome with SNAP disease association dataset for upto four node graphlets. Our approach considers retains the links that participate in the given graphlet while discrading the rest whereas Windels et al. considers adding links between any pair of nodes that are part of the same graphlet. It shows that our approach finds more significant modules. We only show induced networks that are different in the two methods as for G0, G2, and G8, the two methods are equivalent.

| Disease | Best adj. p-value |
|---|---|
| Thrombosis | 0.09 |
| Chronic Myeloid Leukemia | 1.0 |
| Age related macular degeneration | 0.08 |
| Glioblastoma | 1.0 |

Table S1: Table of p-values of $G_0$-based clustering in SNAP and selected DisGeNET associations as shown in Fig. 6 (main text).

| Trait | Category | GWAS name | Reference |
|---|---|---|---|
| Coronary Art. Dis. | Cardiovascular | EUR.ASN.CAD.cad.add.160614.website.txt.tgz | Nikpay et al., *Nat Gen* 2015. |
| Body Mass Index | Anthropometric | EUR.BMI.ENGAGE1000G_BMI.txt.tgz | Horikoshi et al., *PLoS Gen* 2015. |
| Type 2 Diabetes | Glycemic | EUR.DIAGRAMv3.2012DEC17.T2D.txt.gz | Morris et al., *Nat Gen* 2012. |
| Overweight | Anthropometric | EUR.GIANT_OVERWEIGHT_Stage1_ Berndt2013_publicrelease_HapMapCeuFreq.txt.gz | Berndt et al., *Nat Gen* 2013. |
| Alzheimer's Disease | Neurodegenerative | EUR.IGAP_stage_1.txt.gz | Lambert et al., *Nat Gen* 2013. |

Table S2: Table of GWAS traits and names.

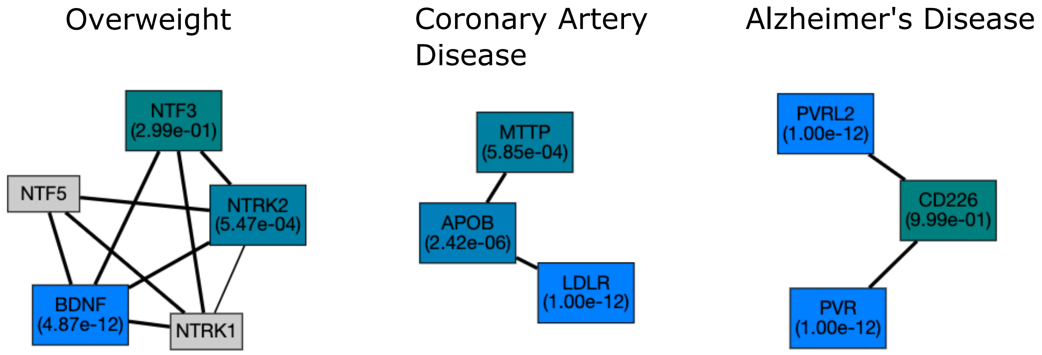


Figure S6: Additional GWAS-trait associated significant modules found by higher-order graphlets that are not found either by $G_0$-based clustering or by one of the top 5 methods from the DREAM challenge. The associated traits Overweight, Coronary Artery Disease, and Alzheimer's Disease are detected by graphlets $G_{24}$, $G_{22}$, and $G_5$ respectively.
